# Tumor clone dynamics in gastro-esophageal cancer organoids reveal a non-genetic memory of neoadjuvant chemotherapy via downregulation of NFκB signaling

**DOI:** 10.1101/2025.07.29.667467

**Authors:** Thi Tuong Vi Dang, Tim Schmäche, Dmitrii Severinov, Vivian Mittné, Juliane Fohgrub, Franziska Baenke, Daniel E. Stange, Anna R. Poetsch

## Abstract

Adenocarcinomas of the gastroesophageal junction exhibit genetic and non-genetic heterogeneity that impact clinical outcomes, though the underlying mechanisms behind drug resistance remain poorly understood. We integrated bulk whole-genome sequencing (WGS) and single cell RNA sequencing (scRNA-seq) data from patient-derived organoid lines generated from drug resistant gastric tumors of three patients before and after chemotherapy with FLOT (5-fluorouracil, leucovorin, oxaliplatin, and docetaxel), investigating both *in vivo* and *ex vivo* treatment effects. We found that inter-patient variability of gene expression exceeds intra-patient differences and predominantly shapes the expression profiles. Integration of WGS-inferred cancer phylogenies with scRNA-seq data allowed us to associate genetic clones with the individual cells’ transcriptional program and to track the genetic and transcriptomic history of dominant genetic clones in post-treatment samples relative to the corresponding primary tumor. Notably, *in vivo* treated samples appeared to be transcriptionally distinct from the untreated counterparts, marked by sustained NF-κB down-regulation, which suggests that they retain an immune-mediated imprint of the prior therapy. Changes in the clonal composition of a tumor alone cannot explain the post-chemotherapy NF-κB-associated transcriptional reprogramming. Instead, non-genetic mechanisms shape the altered transcriptomic landscape, particularly a distinct subpopulation of epithelial cells that specifically express pro-inflammatory cytokines, key components of the NF-κB regulatory network. These observations support a model of transcriptional reprogramming after FLOT treatment, which is most likely independent of genetic evolution and consequently potentially reversible. Downregulated NF-κB signaling may thus represent a candidate pathway change for predictive response assessment and/or NF-κB-stimulating co-therapeutic strategies to overcome FLOT resistance.

## Introduction

Gastric cancer is one of the most deadly cancers globally and represents a major health emergency considering that among a cohort of young people born in 2008–2017, 15.6 million people globally are projected to develop gastric cancer in their lifetime^1^. Multi-target treatments, including the peri-operative FLOT regimen (5-fluorouracil/5-FU, leucovorin, oxaliplatin, docetaxel), have improved outcomes to a median overall survival of 50 months^2^, but long-term survival remains poor, with about 50% of patients experiencing relapse, disease progression or death within two years. Genetic tumor heterogeneity is recognized as a major contributor to treatment resistance and disease recurrence^3^. As cancer cells evolve, they accumulate somatic mutations, some of which can drive tumor progression and confer drug resistance. Yet, it is not clear whether the clones that harbor resistance mutations already exist at the point of treatment, or whether the chemotherapeutic agents, which can be mutagenic, may induce such mutations. In addition, non-genetic tumor heterogeneity and adaptive processes allow for escapes from the treatment and the associated immune response^4^. Notably, non-genetic mechanisms are very dynamic and adaptable, yet may also be reversible. This can be mediated by the tumor microenvironment (TME) or changes in the tumor cells themselves, e.g. their epigenome or transcriptome. To address non-genetic changes in the tumor cells directly, it becomes necessary to model the tumors without their microenvironment. For this purpose, patient-derived organoids (PDOs) are a very suitable model. While they do not include the TME and are therefore independent of it, they encapsulate the clonal heterogeneity of the tumor and facilitate *ex vivo* treatment studies and the investigation of the associated clonal development of the tumor.

As part of the “Outcome prediction of systemic treatment in esophagogastric carcinoma” (Opposite) study (ClinicalTrials.gov #NCT03429816), we are applying *ex vivo* FLOT treatment to PDOs, which allows us to differentiate pathological responders from non-responders^5^. For the three patients investigated in this study, we additionally generated PDOs from resected tumors after neoadjuvant FLOT treatment *in vivo*. This gives us the unique opportunity to study the clonal structure of primary tumors and their dynamics after FLOT treatment both *in vivo* and *ex vivo*.

For this purpose, we performed bulk whole genome sequencing (WGS) and single-cell RNA sequencing (scRNA-seq) on the three organoid lines for each patient, i.e. from i.) the treatment-naïve, ii) the *ex vivo* treated former treatment-naïve organoid, and iii) the *in vivo* treated tumors. While bulk WGS reliably identifies somatic mutations, scRNA-seq profiles provide gene expression at single-cell resolution, which reveals transcriptional intra-tumor heterogeneity. By integrating these two complementary modalities, we resolved tumor clonal composition and simultaneously explored genotype-phenotype relationships at single-cell resolution. Here, we found notable transcriptomic differences dependent on FLOT neoadjuvant chemotherapy, when given *in vivo* and *ex vivo*, specifically a downregulation of the NF-κB pathway in a subpopulation of cells after *in vivo* treatment independent of observed genetic clone dynamics. This highlights a transcriptional imprint of *in vivo* inflammation-driven treatment effects, that is preserved in PDOs. Such a signature of FLOT resistance may enable predictive response assessment and suggests that co-therapeutic strategies targeting NF-κB activation could help overcome resistance.

## Results

To investigate clonal evolution in response to FLOT treatment in adenocarcinomas of the gastro-esophageal junction (GEJ), we performed WGS and scRNA-seq on a total of nine organoid lines, derived from tumor samples from three independent GEJ patients. OO77 and OO100 were classified as GEJ I (lower third of the esophagus), while OO99 was classified as GEJ II (tumor center at the gastroesophageal junction). We obtained organoids established from the treatment-naïve primary tumor, organoids generated from the tumor after *in vivo* neoadjuvant chemotherapy with FLOT, and those where FLOT (5.15 µM 5-FU, 5.35 µM oxaliplatin, 0.6 nM docetaxel, 10 µM calcium folinate) was added directly to the organoid derived from the treatment-naïve biopsy (*ex vivo*) (**Fig1A**). *Ex vivo* treatment was performed for 4 days, followed by a 3-day recovery period. After regrowth (2–4 weeks), the treatment was repeated. Samples were collected after complete recovery from the second cycle and processed for WGS and scRNA-seq. Together with the WGS and scRNA-seq data from the post-treatment derived PDOs, this dataset enables the investigation of how tumor genetic and transcriptomic heterogeneity react to drug treatment, administered either *in vivo* or *ex vivo*. After quality control, scRNA-seq data of nine tumor organoid lines contained gene expression information for a total of 43,366 epithelial cells (median 4,072 cells per sample, 19,592 UMIs and 4,873 genes per cell; **FigS1**).

**Figure 1.**
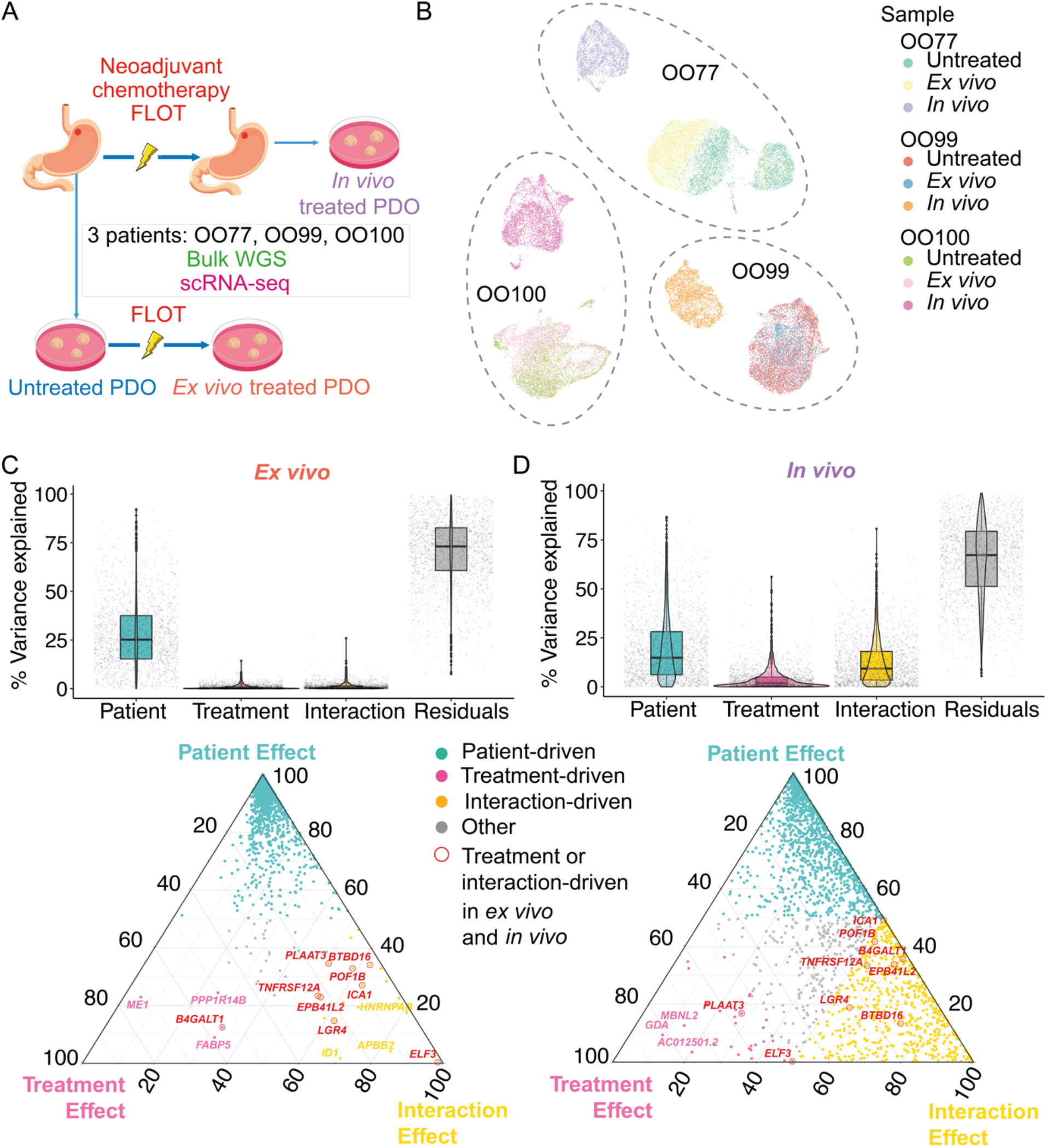
Inter-patient heterogeneity surpasses intra-patient heterogeneity and significantly influences gene expression. (**A**) Schematic of experiment design and data used in the study. FLOT = 5-fluorouracil, leucovorin, oxaliplatin, docetaxel; PDO = Patient Derived Organoid; WGS = Whole Genome Sequencing; scRNA-seq = single cell RNA-seq. (**B**) Uniform Manifold Approximation and Projection (UMAP) embeddings of 43,366 cells from nine samples belonging to three patients. Cells are colored by sample of origin. (**C&D**) Top: Box plot and Violin plot showing percentage of explained variance when fitting a linear model for each gene among the top 2000 most variable genes with covariates including patient, treatment and their interaction effect. Bottom: Ternary plots illustrating the relative contributions of patient, treatment, and their interaction to gene expression variability among the top 2000 most variable genes. Each point represents a gene, colored by the primary source of explained variance: blue if >50% is attributed to the patient, pink for treatment, orange for interaction, and gray if no single factor exceeds 50%. Genes influenced by treatment or interaction in both *ex vivo* (**C**) and *in vivo* (**D**) settings are highlighted with red circles. Selected gene names are labeled.

### Inter-patient heterogeneity exceeds intra-patient heterogeneity

First, we investigated the single-cell transcriptome by identifying highly variable features followed by Uniform Manifold Approximation and Projection (UMAP; **Fig1B**). Cells derived from the same individual are generally clustered together, which reflects stronger inter-patient than intra-patient transcriptional heterogeneity. This observation is consistent with previously reported transcriptome variability across patients in other cancers^6^ and is not surprising given the patient-specific variability of gastric PDOs in morphology and drug response^7^. To dissect the relative contributions of patient identity, treatment, and their interaction, we applied linear models to each of the top 2,000 most variable genes relative to both *ex vivo* and *in vivo* treatment (**Fig 1C & D**). In both contexts, patient identity explains the largest proportion of gene expression variance. Treatment and interaction effects are more pronounced in the *in vivo* condition than in an *ex vivo* setting. Notably, while a large proportion of genes show expression patterns associated with the patient effect or interaction effect of it with either *ex vivo* or *in vivo* treatment, nine genes consistently show association with both *ex vivo* and *in vivo* treatment. These include *ELF3, LGR4*, and *TNFRSF12A*, genes that were previously reported as mediators of chemoresistance to cisplatin and 5-FU through pathways such as epithelial-mesenchymal transition (EMT)^8^, Wnt/β-catenin signaling^9^, and NF-κB signaling^10^, respectively, albeit in the context of an upregulation in former studies, whereas we observe a downregulation. Our results suggest that inter-patient heterogeneity dominates transcriptional variation. When chemotherapy is applied both *in vivo* and *ex vivo*, it affects key cancer-related pathways involved in the treatment response, yet *in vivo* treatment induces additional effects not captured *ex vivo*.

### *In vivo* treated samples are transcriptionally distinct from untreated samples and exhibit reduced NF-κB signalling

We next investigated in greater detail how treatment alters tumor gene expression, aiming to understand the molecular changes induced by neoadjuvant therapy and *ex vivo* drug exposure. Indeed, *in vivo* treated samples, in comparison to *ex vivo* treated PDOs, are transcriptionally more distinct from untreated samples (**Fig2A**). This may be due to the clonal origin of the organoids, which is not necessarily shared between the untreated biopsy and the resected tumor after *in vivo* treatment. Focusing on the transcriptomic changes, the majority of genes (24.68%, 9.45%, and 10.25%, for OO77, OO99, and OO100, respectively) are deregulated in response to either *in vivo* or *ex vivo* treatment. Only a small subset (2%, 0.22%, and 0.26% for OO77, OO99, and OO100, respectively) of genes showed similar expression trends when comparing *in vivo* or *ex vivo* treated samples to the matched untreated samples (**Fig S2**). Comparing the effects of *in vivo* and *ex vivo* treatment, more genes are differentially regulated between *in vivo* treated samples and the corresponding untreated control with n=7298 vs 951, 2839 vs 133, and 3069 vs 159, for OO77, OO99, and OO100, respectively (**Fig S2**). In conclusion, *ex vivo* treatment of the PDOs only captures a subset of the effect we observe from *in vivo* treatment.

**Figure 2.**
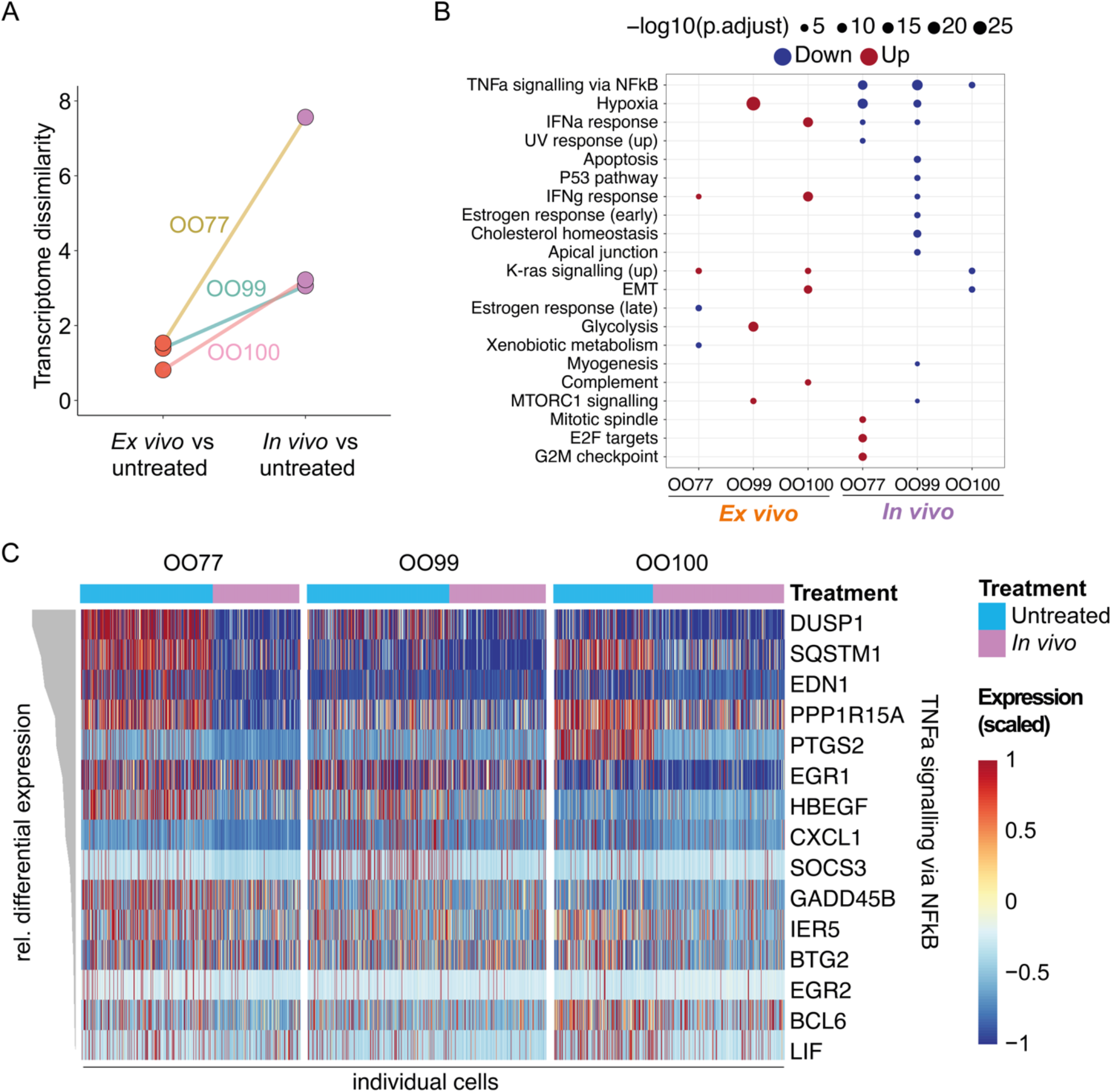
*In vivo* treated samples are transcriptionally distinct from untreated samples and exhibit downregulation of the NF-κB signaling pathway. (**A**) Transcriptome dissimilarity between post-treatment samples and the corresponding untreated samples of three patients OO77, OO99, and OO100. (**B**) MSigDB HALLMARK gene sets that are significantly enriched (Benjamini-Hochberg adjusted p < 0.01) using the significant genes identified from pseudobulk differential analysis between post-treatment samples and the corresponding untreated samples. (**C**) Heatmap showing scaled expression of genes from the HALLMARK_TNFA_SIGNALING_VIA_NFKB gene set that are consistently downregulated across all three patients with a log2fc > 0.5 following *in vivo* treatment. Rows represent genes and columns represent cells, sorted by relative differential expression as the mean scaled logcount expression of untreated minus *in vivo* treatment (depicted in grey).

We next interrogated which gene pathways are targets of FLOT induced deregulation. Combining the scRNA-seq data to a pseudo-bulk differential gene expression analysis, the gene set “TNFa signaling via NFκB” was consistently downregulated in *in vivo* samples across all three patients, whereas such changes were not observed in *ex vivo* samples. This implies a reduction of NF-κB signaling after *in vivo* treatment (**Fig2B**). Indeed, expression levels of multiple genes functioning in NF-κB signaling were reduced in *in vivo* compared to untreated samples (**Fig2C**), although some cellular heterogeneity remained.

### Bulk WGS-derived tumor evolution trees reveal genetic difference between untreated and post-treatment samples

Transcriptional changes between pre- and post-treatment samples may reflect underlying genetic shifts and heterogeneity. Therefore, we reconstructed phylogenetic trees using all three bulk WGS samples of each patient (**Fig3&FigS3**) and deconvoluted the mutagenesis history in each patient (**Fig S3**). Importantly, the phylogenetic trees allow the timing of key driver mutations and structural aberrations, based on the clonality and abundance level of these genetic aberrations. *TP53* is mutated in OO77 and OO99, and its inhibitor, E3 ubiquitin-protein ligase (*MDM2*), is amplified in patient OO100. These events occurred early in tumor development beside other cancer driver mutations typical for gastric cancer^49^. Clonal composition shifts dramatically after treatment, both *in vivo* and *ex vivo*, which illustrates the differences in tumor genetic composition in pre- and post-treatment samples. While, for OO77, the ancestral clone for both the *in vivo* and *ex vivo* treated samples is the general common ancestor clone 1, for OO99 and OO100 the *in vivo* treated resistant clone arises from clone 2, which is shared with the untreated sample, but does not survive the clonal sweep of *ex vivo* treatment. It is noteworthy that by comparing the number of differentiated clones (four vs one in OO77, three vs two OO99 and three vs three in OO100), the clonal reconstruction for *ex vivo* treated samples is more differentiated than for *in vivo* treated samples. This may be due to the better resolution in the untreated sample providing the direct ancestor clones of the *ex vivo* treated sample, whereas the *in vivo* treated sample may be derived from a separate clone combination that we cannot deconvolute in similar resolution. Interestingly, despite the surprisingly light effects of the *ex vivo* treatment on the PDO’s transcriptomes, clonal dynamics go through a substantial bottleneck. In all three samples, the majority of the clones present in the untreated sample do not survive the *ex vivo* treatment and smaller clones persist and evolve. In OO99 and OO100, the shared ancestor clone for both treatments disappears *ex vivo*. Together, this suggests that the mechanisms that drive clonal dynamics *in vivo* and *ex vivo* are disconnected or have a very strong stochastic component, which leads to non-concordant changes of diverse cell populations, when characterized both genetically and non-genetically. Yet these analyses were performed separately for each modality, which may hide concordant behaviors.

**Figure 3.**
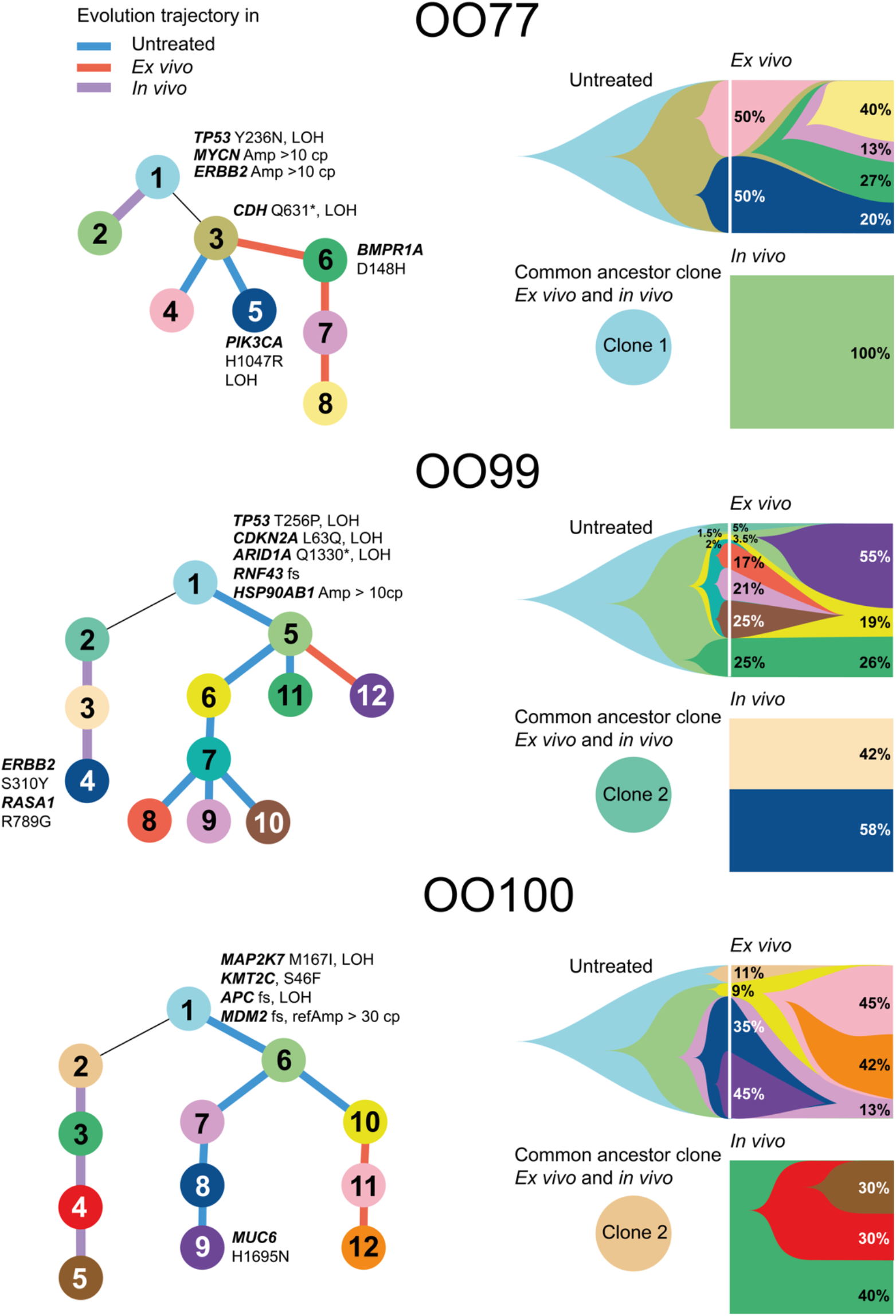
Bulk WGS-derived tumor evolutionary trees reveal genetic differences between pre and post-treatment samples. **Left**: Phylogenetic trees derived from bulk WGS data annotated with putative driver mutations and Copy Number Variants (CNVs) of known gastric cancer driver genes. Each circle is a clone as a group of mutations which have similar clonality and cancer cell fraction. Colored lines represent the sample in which the clone evolved, if it can be uniquely assigned. **Right**: Reconstruction of the clonal proportions and their evolution. The white line depicts the timepoint of the untreated sample from which the evolution from *ex vivo* treatment starts. The common ancestor clone depicts the clone that is present in the untreated sample and the *in vivo* treated sample. Given that the *in vivo* treated sample can be derived from a different location, this clone represents the earliest commonality between all samples - even if it did not persist during *ex vivo* treatment. Clonal reconstruction at endpoints is quantitative and marked with the associated proportions, yet clonal dependencies and dynamics in between are qualitative.

### scRNA-seq allows to refine clonal reconstruction and find resistant clones in treatment naïve samples

To achieve a refined resolution of the clonal reconstruction, we performed clone labelling for each cell in the scRNA-seq data based on the mutations that can be detected in the transcriptome. The clone composition observed in scRNA-seq data generally aligns with that derived from bulk WGS, yet in OO77 it became apparent that the clone assignment is incomplete (**FigS4**). From scRNA-seq data, clone 4 cannot be verified and instead appears to be a subset of clone 5, which in turn shows the identified *PIK3CA* mutation only in a subclone, subsequently called clone 5b **(Fig5 & FigS4)**. In this refined clone reconstruction of OO77, it becomes clear that the *in vivo* resistant clone 2 is both genetically and transcriptionally distinct (**Fig4A & FigS4**). The emerging clone 6 in the *ex vivo* treated sample is both genetically and transcriptionally largely distinct from the two variants of clone 5, where clone 5a shows an overlapping set of mutations with clone 6 and can therefore be considered its ancestor. In both OO99 (**Fig4B**) and OO100 (**Fig4C**), a large proportion of the single cells of the untreated and *ex vivo* treated sample cannot be assigned with certainty to a genetic clone, which skews the percentage of recovered clones. However, from the recovered clones, it becomes clear that genetic heterogeneity cannot explain the transcriptional heterogeneity in these samples. In both OO99 and OO100, this is different for the clones that emerge in the *in vivo* treated sample that are both genetically distinct and occupy a separate territory with its single cell transcriptomes. In this refined reconstruction the sparsity of scRNA-seq data prevents us from assigning every cell to a clone. However, recurrent discovery of mutations in the scRNA-seq data allows tracing of specific clones in single cell resolution. In OO100, this manifests that mutations assigned to clones 3 and 4, the resistant clones from *in vivo* treatment, are detectable already in 47 and 7 untreated cells as well as in 35 and 10 *ex vivo* treated cells, respectively. While this was not detectable from the clonal reconstruction using bulk WGS data, it indicates that the resistant clones were already present at the time of *in vivo* treatment and then selected. Yet, transcriptionally, these cells do not group with the *in vivo* treated samples, which suggests that we are either missing additional genetic changes that would define a separate clone, or the distinct transcriptomic separation of the PDOs from the resistant clones is independent of their genetic makeup. In summary, the treatment-related clones in OO77 are both genetically and transcriptionally distinct, while in OO99 and OO100 only the *in vivo* treated sample has a distinct transcriptome. Although the *ex vivo* samples also genetically evolve, the changes do not result in the evolving clones being transcriptionally distinct. In addition, for OO100, the integrated clonal reconstruction reveals the pre-existent resistant clone that dominates after *in vivo* treatment to be present also in the untreated and *ex vivo* samples, but without the transcriptional distinction that characterises the same clone in the *in vivo* treated sample.

**Figure 4.**
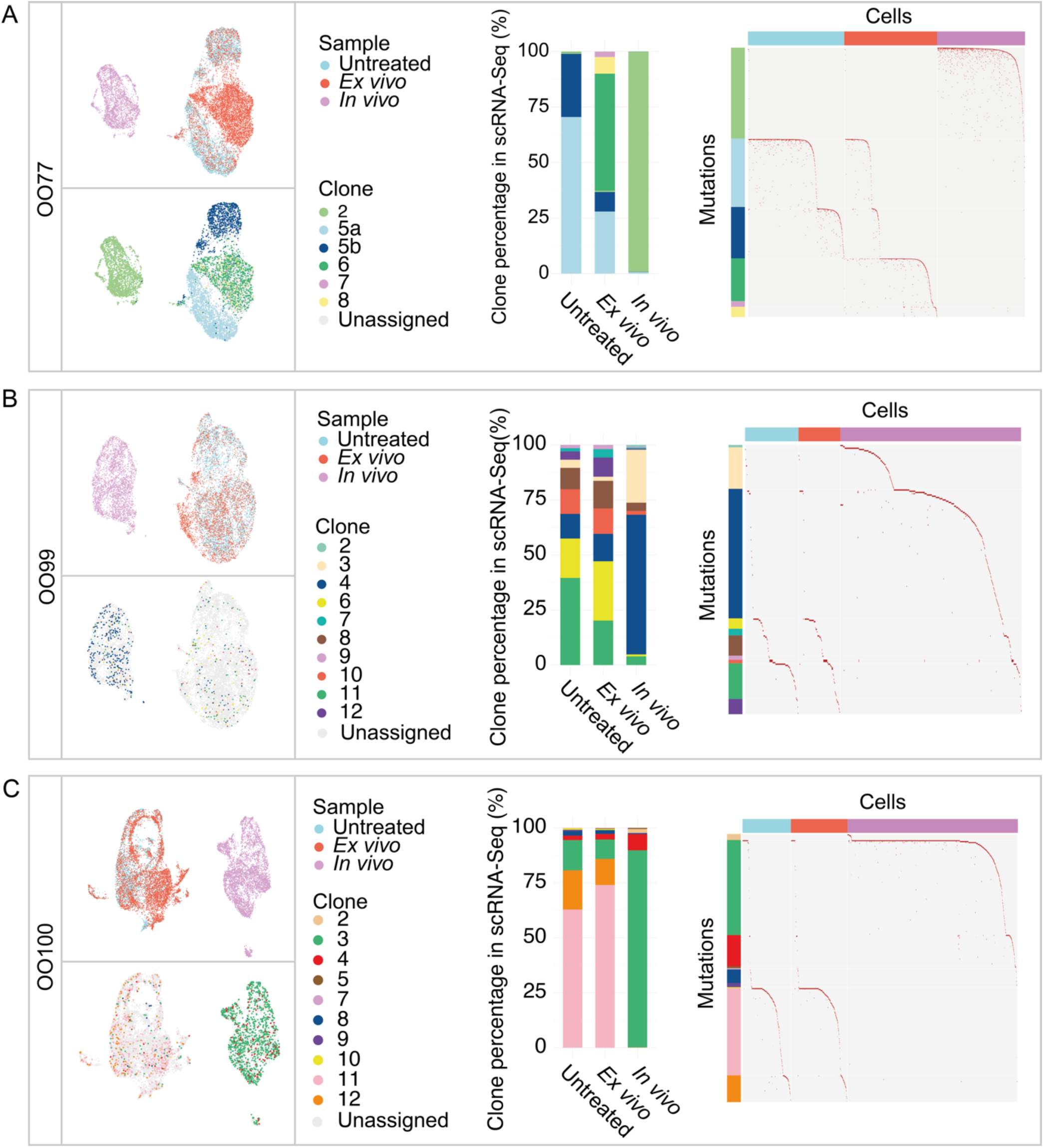
Refinement of clonal reconstruction to find resistant clones in treatment naïve samples. (**A-C**) Mapping clones from phylogenetic trees to scRNA-seq data in OO77, OO99, and OO100, respectively. **Top-left**: UMAP embeddings of cells from all three samples untreated, *ex vivo, in vivo*. Cells are colored by samples of origin. **Bottom-left**: The same UMAP embeddings as in the top panels. Cells are colored by assigned clones. **Center:** Stacked bar plots showing the percentage of each clone among labeled cells in each sample according to scRNA-seq data. **Right**: Mutation matrix of clones relative to treatment. Rows represent mutations from bulk WGS-derived phylogenetic trees, annotated according to their clone membership within the trees. Columns correspond to cells in scRNA-seq data differentiated by treatment. Rows are sorted by clone and mutation recurrence, columns by treatment and the presence of mutation.

**Figure 5.**
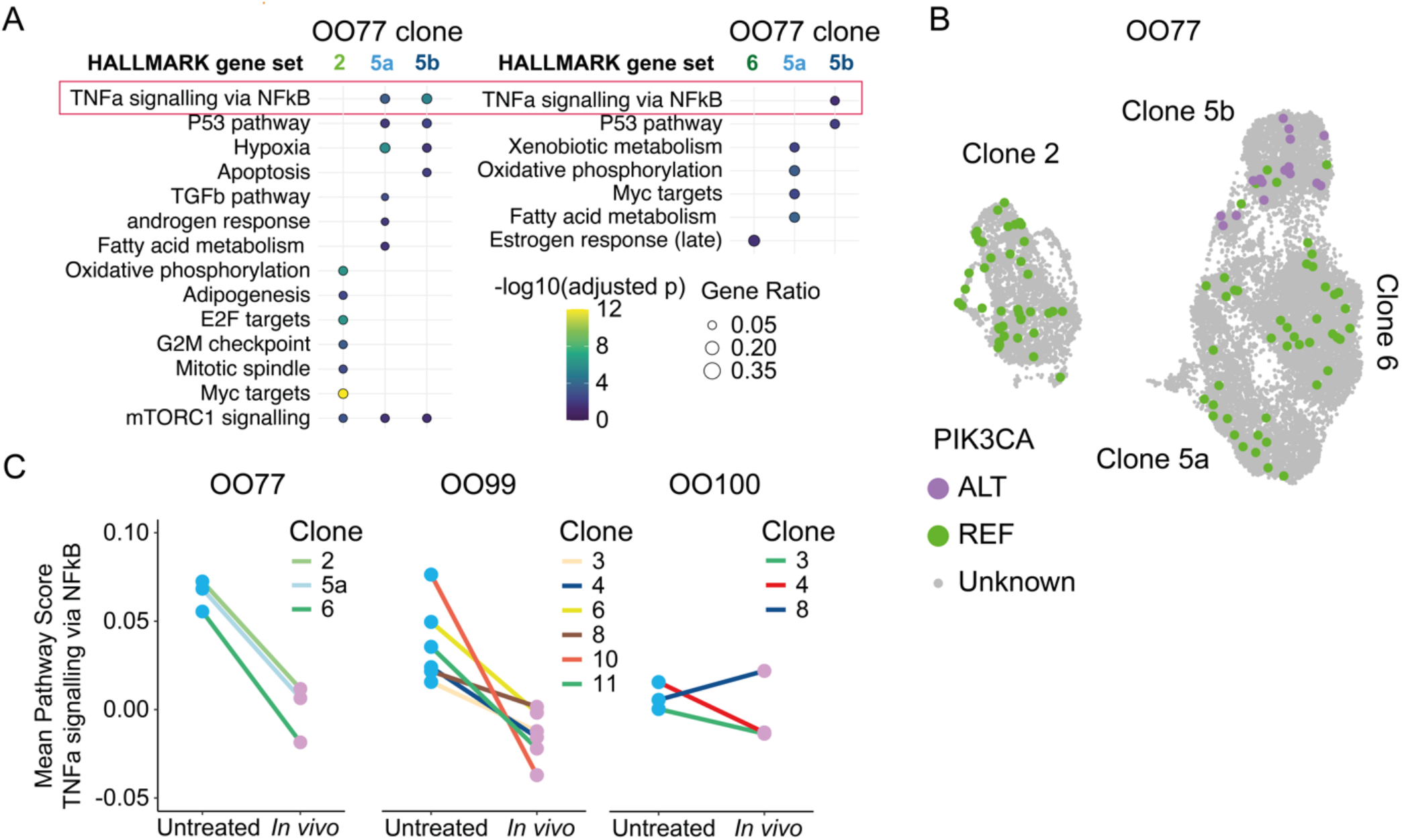
The down-regulation of the NF-κB pathway following *in vivo* treatment is not fully attributed to genetic factors but reflects a global transcriptional shift. (**A**) Gene expression properties of genetic clones in patient OO77, where clone 2 is the clone that emerges from *in vivo* treatment and clone 6 emerges from *ex vivo* treatment. MSigDB HALLMARK gene sets that are significantly enriched (Benjamini-Hochberg adjusted p-value < 0.01) among the marker genes of each genetic clone of OO77. (**B**) UMAP representation of cells in OO77 which are detected with either reference (REF) or mutant (ALT) allele of the mutation *PIK3CA* p.H1047R. The same UMAP coordinates as in Fig.4 for this patient were used. (**C**) Genetic clone-specific comparison of HALLMARK_TNFA_SIGNALING_VIA_NFKB gene set between untreated and *in vivo* treated samples in three patients.

### Following *in vivo* treatment, the NF-κB pathway is downregulated within genetic clones

We next asked whether the NF-κB pathway downregulation observed in *in vivo* treated samples (**Fig2**) could be explained by shifts in the underlying clonal composition of tumor cells. In OO77, the clones 5a/b, present in the untreated sample, exhibited higher NF-κB activity compared to clones 2 and 6, which thrive after *in vivo* and *ex vivo* treatment, respectively (**Fig5A**). In the case of clone 5b, this can be explained by the gain-of-function mutation in *PIK3CA* (**Fig5B**), which is known to activate NF-κB signaling^11^. This suggests that chemotherapy may either preferentially eliminate clones with high baseline NF-κB signaling or give those with lower NF-κB phenotype a growth advantage. To investigate this relationship in more detail, we compared NF-κB activity within each clone before and after treatment (**Fig5C**). We observed that NF-κB signaling decreases after treatment in nearly all clones. The only exception was clone 8 in OO100, which is represented by a limited number of cells (n = 4) in the *in vivo* treated sample. Taken together, these results indicate that while clonal selection might contribute to the observed reduction in NF-κB signaling, it cannot fully explain the extent of pathway suppression.

### Down-regulation of the NF-κB pathway following *in vivo* treatment reflects a general transcriptional shift towards reduced cytokine expression

Due to the lack of clear links to genetic sub-clonality, we explored transcriptional heterogeneity and integrated all 43,366 cells across conditions to perform unsupervised clustering. We identified nine cell clusters, which are shared across all samples (**Fig6A** and **Fig S6A&B**). Among the non-cycling cells, cluster 9 shows elevated expression of NF-κB pathway activity (**Fig6B** and **Fig S6C-E**). However, even within each transcriptional cluster, NF-κB activity declines following treatment (**Fig6C**), with cluster 9 showing the highest baseline expression. These findings suggest that the downregulation of NF-κB signaling following *in vivo* chemotherapy is not solely a result from the selection of pre-existing clones or cell states with static phenotypes, but also reflects broader transcriptional reprogramming.

**Figure 6.**
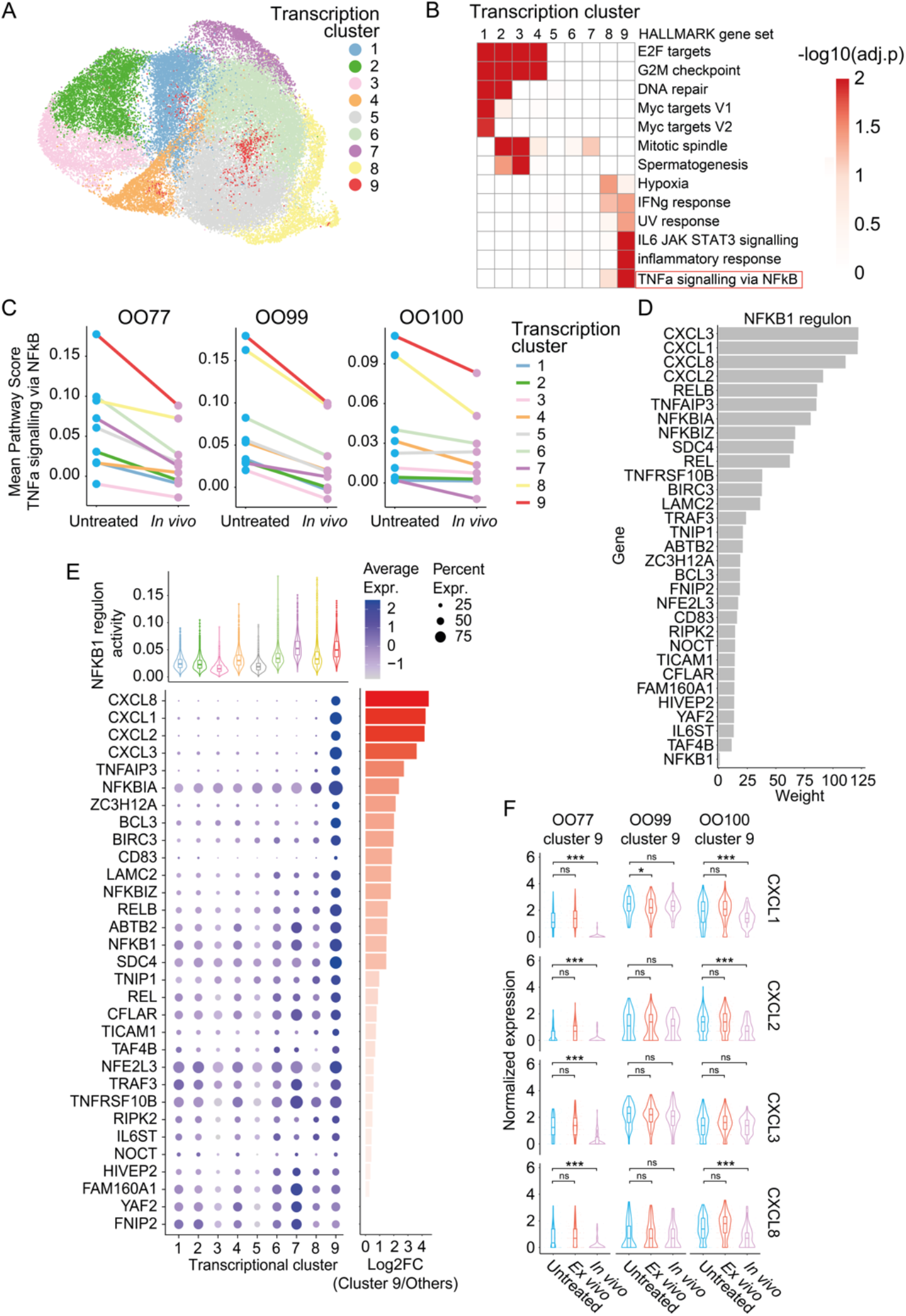
Tumor-wide NF-κB pathway suppression after *in vivo* treatment can be pinpointed to an inflammatory epithelial cell cluster. (**A**) UMAP representation of transcriptional clusters shared across nine samples after CCA-based integration. (**B**) Heatmap showing MSigDB HALLMARK gene sets that are significantly enriched (BH adjusted p-value < 0.01) using the marker genes of each transcription cluster. (**C**) Intra-transcriptional cluster comparison of HALLMARK_TNFA_SIGNALING_VIA_NFKB gene set between untreated and *in vivo* treated samples in three patients. (**D**) Barplots showing genes belonging to NFKB1 regulon with their weights as inferred by SCENIC. (**E**) Top: Violin plots and boxplots showing NFKB1 regulon activities across cells belonging to transcriptional clusters. The middle line represents the median, the box denotes the interquartile range (IQR), and the whiskers extend to 1.5 times the IQR. Bottom left: Dotplots showing gene expression levels across transcriptional clusters of marker genes of cluster 9. Bottom right: Differential expression between Cluster 9 versus the other clusters. (**F**) Violin plots and boxplots showing gene expression levels of cytokines in pre- and post-treatment. The middle line represents the median, the box denotes the interquartile range (IQR), and the whiskers extend to 1.5 times the IQR. Statistical significance was assessed using the Student’s t-test, with significance levels indicated as follows: p-value > 0.05 (ns), p-value < 0.05 (*), p-value < 0.01 (**), p-value <0.001 (***).

Given the dynamic regulation of NF-κB signaling across the samples, we next investigated the transcriptional mechanisms underlying this pathway’s activity. We reconstructed the NF-κB signaling gene-regulatory network using the Python implementation of SCENIC (Single-Cell rEgulatory Network Inference and Clustering)^12^, pySCENIC v0.12.1^46^, and identified the *NFKB1* regulon in our samples, which is primarily driven by pro-inflammatory cytokines such as *CXCL1, CXCL2, CXCL3*, and *CXCL8* (**Fig6D**). Consistent with the results above, transcriptional cluster 9 highly expresses these cytokines along with other NF-κB pathway components, which include *RELB, NFKBIA*, and *IER3* (**Fig6E**). *In vivo* treated samples showed a marked reduction in *CXCL1, CXCL2, CXCL3*, and *CXCL8* expression in cluster 9, most notably in OO77, and to a lesser extent in OO99 (**Fig6F**). This shows that a subpopulation of the organoids have high NF-κB signaling with the expression of the associated cytokines, despite the absence of an immune system when culturing them. Interestingly, while *ex vivo* treatment does not affect NF-κB signaling that cannot be explained by genetic clonal dynamics with *PIK3CA* mutations, *in vivo* treatment alters the proinflammatory cell population without detectable genetic changes. This suggests that a transcriptional memory of the TME is conserved in the PDO culture and, a reduction of NF-κB signaling could be as an immune escape mechanism induced by *in vivo* FLOT treatment.

## Discussion

In this study we used PDOs from three patients to investigate the clonal and single cell dynamics of GEJ in response to *in vivo* and *ex vivo* FLOT treatment. We found that inter-individual variability has a greater effect on single-cell transcriptomes than treatment. *In vivo* treatment shows larger impact on single cell transcriptomes than *ex vivo* treatment, despite both showing genetic changes in clone composition. Unlike the *ex vivo* treated PDOs, *in vivo* treatment involves tumor interaction with the immune system and associated selective processes. In all three patients we observe a downregulation of NF-κB signaling, independent of genetic changes, most prominently in a coherent cluster of cells with consistently high NF-κB signaling and expression of associated cytokines.

Assigning genetic clones to single cells showed that such treatment-adaptations are likely not anchored in genetic changes, but rather represent a transcriptional memory. The stability of this memory is striking, given that the interaction with the immune system would have taken place before the tumor was used to generate organoids, yet the transcriptional program persists even after *ex vivo* cultivation.

Also, the direction of change in NF-κB signaling is noteworthy. The same pathway is generally regarded as being upregulated after chemotherapy, which leads to survival signaling and associated chemotherapy resistance^13–15^. This raises the question why a downregulation would be protective. NF-κB signaling promotes immune system recruitment by increasing the expression of pro-inflammatory cytokines, which can exert anti-oncogenic effects^16^. Its downregulation may therefore represent a mechanism of immune evasion. FLOT includes a platinum-based compound whose clinical effect is partly mediated through interactions with the immune system^17^. Immune evasion would therefore be a strategy to reduce treatment effects. The fact that this adaptation is achieved through transcriptomic reprogramming, rather than genetic selection, demonstrates the plasticity of resistance mechanisms and hints that intercellular communication within the tumor enables such changes to transcend clonal boundaries. This finding also represents an opportunity for clinical prognosis and intervention. The fact that PDOs retain the memory of their FLOT resistance in the form of NF-κB signaling activity may provide avenues in using NF-κB signaling as a predictive marker for FLOT resistance. In addition, since the resistance mechanism is mediated by gene regulatory networks rather than genetic changes, it may in principle be reversible. If the down-regulation of NF-κB signaling indeed represents a mechanism of resistance through immune evasion, stimulation of the pathway through cytokines or inducing local inflammation have the potential for sensitization of tumors and enhance the immune response.

In conclusion, careful dissection of clonal dynamics and single cell transcriptomics in gastric cancer organoids has revealed a downregulation of NF-κB signaling via transcriptional memory in organoids. Downregulation of this pathway is associated with resistance to FLOT in GEJ, which has the potential to be exploited for predictive and therapeutic use.

## Methods

### Sample collection

Samples from treatment-naïve patients were collected as previously described^5^. Briefly, endoscopic biopsies (2–3 per patient) were obtained from the primary tumor site prior to neoadjuvant FLOT chemotherapy. Tissues were transported in ice-cold NaCl and processed immediately for PDO generation. Matched post-treatment samples were obtained from surgical resection specimens after neoadjuvant FLOT chemotherapy and kept in cold PBS until processing. Ethical approval was obtained for all procedures (EK 169052018, BO-EK-421092022).

### PDO generation and culture

PDOs were established and maintained as previously described^7^. Tumor tissue was enzymatically digested using dispase II and collagenase XI. The resulting cell suspension was washed and embedded in Matrigel for cultivation in gastric organoid medium (detailed composition remains unchanged). Y-27632 (10 µM) was added during early passages to enhance survival. Organoids were passaged 1–2 times per week depending on growth characteristics.

### *Ex vivo* FLOT treatment

PDOs were treated with a FLOT mixture (5.15 µM 5-FU, 5.35 µM oxaliplatin, 0.6 nM docetaxel, 10 µM calcium folinate) for 4 days. After treatment, the medium was replaced with a standard cultivation medium for 3 days. Organoids were then dissociated mechanically and cultured for 2-4 weeks until they reached sufficient growth before being treated a 2^nd^ time. Samples used for downstream analysis were harvested only after full recovery from the second treatment cycle.

### Whole genome sequencing (WGS)

WGS was performed on DNA extracted from PDOs. 100 ng of DNA per sample was used for library preparation using the TruSeq DNA Nano kit (Illumina), following the manufacturer’s instructions. Sequencing was carried out on a NovaSeq X system with paired-end 2 × 151 bp reads, which yields an average genome coverage of 30×. The mean Q30 score across samples was 94.19%.

### Single-cell dissociation and RNA sequencing

Organoids were harvested 48 hours after last passaging, centrifuged (5 min, 300×g), and incubated in Corning Recovery Solution on ice for 30 minutes to remove Matrigel. After washing, they were dissociated into single cells using TrypLE™ Express Enzyme at 37 °C with periodic pipetting. Digestion was stopped using 1% BSA in PBS. Cell viability (>90%) was confirmed via trypan blue staining before proceeding with scRNA-seq. Single cells were processed with the Chromium Next GEM Single Cell 3’ kit (v3.1, dual Index, 10x Genomics) following the manufacturer’s protocol. Libraries were prepped with the FA Quantification NGS Kits (S, SV) and sequencing performed with the NovaSeq 6000 (Illumina). A total of 43,366 epithelial cells were successfully sequenced (median 4,072 cells per sample, 19,592 UMIs and 4,873 genes per cell).

### scRNA-seq data alignment and quality control

Raw single-cell RNA sequencing data were aligned to the human reference genome (GRCh38/hg38) using Cell Ranger v7.1.0 (10x Genomics) to generate unique molecular identifier (UMI) count matrices per gene for each cell. Cells were retained if they expressed at least 200 genes, and genes were included if detected in at least three cells. Further quality control and downstream analysis were performed using Seurat v5^18^. Cells were also excluded if they failed to meet any of the following criteria: UMI counts at least 3,000, detection of at least 500 genes, mitochondrial transcript content lower than 15%, and a transcriptome complexity score (log10[UMI/gene]) of at least 0.8. Potential doublets were identified and removed using DoubletFinder v2.0^19^, applying the recommended multiplexing rate for the 10x Genomics Single Cell 3’ Gene Expression v3.1 chemistry.

### scRNA-seq downstream analysis

We normalized and scaled the filtered UMI count matrix using the function SCTransform v1^48^ in Seurat v5^20^. The top 3,000 most variable genes were used to perform principal component analysis (PCA) with the function runPCA. Louvain graph-based clustering was then performed on the first 30 principal components, and the resulting clusters were visualized using UMAP with the function runUMAP.

To analyze cell states across samples, all cells from nine samples were integrated using the Canonical Correlated Analysis-based approach provided in Seurat using the three untreated samples as reference. Cells were clustered in Seurat using the FindClusters function with the Louvain algorithm at a resolution of 0.35. Cluster-specific conserved marker genes were identified using the Wilcoxon rank-sum test, applying the following criteria: a mean log2 fold change > 0.5 compared to other clusters, expression in > 50% of cells within the cluster, and an adjusted p-value < 0.05. Gene ontology (GO) term analysis was performed using the MSigDB version 7.5.1 on cluster marker genes. Gene set scoring of single cells was performed using the AddModuleScore function in Seurat. For each sample, read counts were aggregated across genes, and transcriptomic dissimilarity scores were computed as one minus the Spearman correlation between sample pairs.

### Linear model fitting

To estimate how much variance in gene expression is explained by different factors, linear models were applied. Since linear models assume that observed expression is an additive combination of covariate effects and random noise, SCTransform normalization is not suitable as it outputs Pearson residuals derived from a regularized negative binomial model, which alters the original expression values and disrupts the additivity required by linear models. Therefore, we normalized the raw single cell RNA-seq count data by library size using the calcNormFactors function from the edgeR package v3.42.4^21^, followed by variance modeling with the voom function from the limma package v3.58.1^22^. We fitted a linear model for each of the top 2,000 most variable genes using the fitExtractVarPartModel function from the variancePartition package v1.32.5^23^. The models included patient identity, treatment, and their interaction as covariates.

### Bulk WGS processing, alignment, somatic SNV and CNV calling and annotation

Sequencing reads were aligned to the reference genome (GRCh38/hg38) using the Burrows-Wheeler Aligner (BWA-MEM) through the nf-core pipeline Sarek v3.2.1. For each patient, aligned reads from all associated PDOs and matched germline control (from blood) were jointly used to call somatic single nucleotide variants (SNVs) and insertions/deletions (indels) using the Mutect2 algorithm from GATK v4.2.0.0. Functional annotation was performed using Ensembl Variant Effect Predictor (VEP v102.0)^24^ and ANNOVAR v06/2020^25^.

Somatic CNV calling for all cancer samples using matched normal samples were performed using AscatNGS v4.5.0^26^. Tumor purity and ploidy values were also estimated by AscatNGS. Called CNV were annotated with ClassifyCNV v1.1.1^27^.

### Phylogenetic tree building

Bulk WGS-derived phylogenetic trees were built using CONIPHER v2.1.0^28^. From the results of Mutect2 and AscatNGS, we implemented an in-house script to prepare input files for CONIPHER which contains somatic SNVs with their estimated average copy numbers in each sample. For OO99 two technical replicates for the untreated sample were used as independent samples. For clusterization and the tree building steps, we used default parameters except setting the parameter minimum_cluster_size to 30. Clusters with a cancer cell fraction (CCF) < 0.15 across all samples were also excluded. If two clusters were found to be in a direct lineage relationship within a single sample and had a CCF difference of less than 0.15, they were merged.

### Mutational Signature Calling

For each patient, the mutational signatures were analyzed using SigProfilerAssignment v0.1.4^29^ to assign previously characterized mutational signatures based on the COSMIC v3.4 reference database. To minimize noise, mutations with a probability ≥ 0.5 of being attributed to known artifact signatures (SBS27, SBS43–SBS60, SBS95) were removed from the phylogenetic trees and downstream analysis. The filtered mutation sets in each clone in the phylogenetic trees were then used as input for SigProfilerExtractor (v1.1.23)^30^ to perform *de novo* extraction of single base substitution (SBS) mutational signatures and all downstream analysis.

Tumor clone composition and dynamics were visualized using the R packages cloneMap v1.0.0^31^, the fishplot version 0.5.2^32^. Fishplots were remade with Adobe Illustrator due to graphical artefacts.

### Putative driver mutation identification

We implemented a pipeline which combined the result of four driver mutation prediction algorithms: MutaGene^33^, CHASMplus^34^, CScape^35^, and CanDra^36^. We only considered mutations that satisfy all three of these criteria as putative driver mutations: i.) Predicted as driver mutations in at least two out of four algorithms; ii) located within genes identified as cancer driver genes in at least two of these databases: COSMIC^37^, INTOGEN^38^, OncoKB^39^, CancerMine^40^; iii) located within genes listed as stomach cancer driver genes as in the multiancestry study by *Totoki et al*^41^.

### Mapping genetic clones from phylogenetic trees to scRNA-seq data and validation

To find somatic mutations detected in bulk whole-genome sequencing (WGS) in single-cell RNA sequencing (scRNA-seq) data, we employed SCReadCounts v1.4.1.25^42^ and Scomatic v1.0^43^. Both tools were used to identify the presence of mutations in individual cells. The results from SCReadCounts and Scomatic were then combined to increase confidence in mutation detection. Clonal assignments were performed based on previously constructed bulk WGS-derived phylogenetic trees for each patient. The following principles guided the assignment of individual cells to specific clones: Only clones inferred from the bulk WGS data that are predicted to still be present at the time of sampling for any of the three samples were considered for cell assignment. For each cell, we assessed the number of detected mutations associated with each clone. A cell was assigned to the clone for which it harbored the largest number of mutations. However, if any mutation associated with a descendant clone was also detected, the cell was instead assigned to that descendant clone.

Clone assignments at the single-cell level were validated by evaluating mutation co-occurrence and clone purity defined as the percentage of cells lacking mutations assigned to other branches (**Fig S4, FigS6**). In patients OO99 and OO100, clone purity was high across all identified clones (except clone 2 of OO99 but this is because only 4 cells were assigned to this clone), supporting the accuracy of the bulk WGS-derived phylogenetic trees in these two patients. However, in patient OO77, initial clone assignment revealed low purity in clone 4, with a substantial fraction of cells containing mutations of clone 5 **(FigS4)**. This suggested a misclassification in the original bulk WGS-inferred tree, likely due to incorrect mutation assignments between clones 4 and 5. To resolve this, we merged the mutations of clone 4 into clone 5 and re-evaluated the resulting clone structure. For each mutation, we computed the centroid of all cells in which it was detected, serving as a proxy for its location in transcriptional space. We then assessed the distribution of pairwise distances between these mutation centroids within the merged clone. The observed bimodal pattern indicated the presence of two transcriptionally distinct subgroups. Using k-means clustering (k = 2) on the mutation centroids, we separated them into two subclones, designated 5a and 5b (**Fig S5**). With this revised classification, clone purity was high across all clones, with each clone containing only mutations assigned to that clone or its ancestors.

Heatmaps were visually enhanced using a gaussian smoothing strategy.

### Pseudobulk Differential Expression and Gene Set Over-representation Analysis

Raw scRNA-seq count matrices were aggregated into pseudobulk samples by summing gene counts within each transcriptomic cluster, treating each cluster as a biological replicate. Differential expression analysis between post-treatment and pre-treatment samples was performed using DESeq2 v1.40.2^44^ on these pseudobulk data. Genes were considered differentially expressed if they had an adjusted p-value < 0.05 and an absolute log2 fold change > 0.5. Gene set over-representation analysis was then conducted using MSigDB v7.5.1^45^ on the differentially expressed genes to identify enriched pathways.

### Gene regulatory network analysis

Gene regulatory network analysis was performed using the Python implementation of SCENIC^12^, pySCENIC v0.12.1^46^, to infer transcription factor (TF)-target gene interactions and to estimate regulon activity at the single-cell level. First, co-expression relationships between TFs and genes were identified using the normalized expression matrix of scRNA-seq data and a curated list of human transcription factors provided by SCENIC authors. Next, motif enrichment analysis was performed using hg38-specific ranking and scoring databases from SCENIC to refine these co-expression modules. Regulons, defined as TFs and their direct target genes, were confirmed by the presence of enriched TF-binding motifs within ±10 kb of each target gene’s transcription start site. For example, the *NFKB1* regulon included genes with the motif “transfac_public M00208” from the TRANSFAC database^47^. Regulon activity per cell was quantified by pySCENIC based on the expression of each regulon’s target genes within each cell.

## Acknowledgements

**ARP** was supported by the Mildred Scheel Early Career Center Dresden P2, funded by the German Cancer Aid. The experimental part was supported by the local funding agency *SMWK [no*. 100615933] under the frame of ERA PerMed.

We thank the whole Biomedical Genomics group at TU Dresden for fruitful discussions and feedback.

## Author contributions

**ARP** conceptualized the computational design of the study, **DES, TS**, and **FB** the experimental design. **FB, DES**, and **ARP** secured funding. **TS** and **JF** generated and cultivated PDOs. **TS** performed *ex vivo* treatment and sample preparation. **FB** and **VM** supported sample preparation, logistics, and communication. **TTVD** and **DS** analyzed the data with input from **ARP. ARP** and **TTVD** wrote the manuscript with input from all authors.

---

**Supplementary Figure 1.**
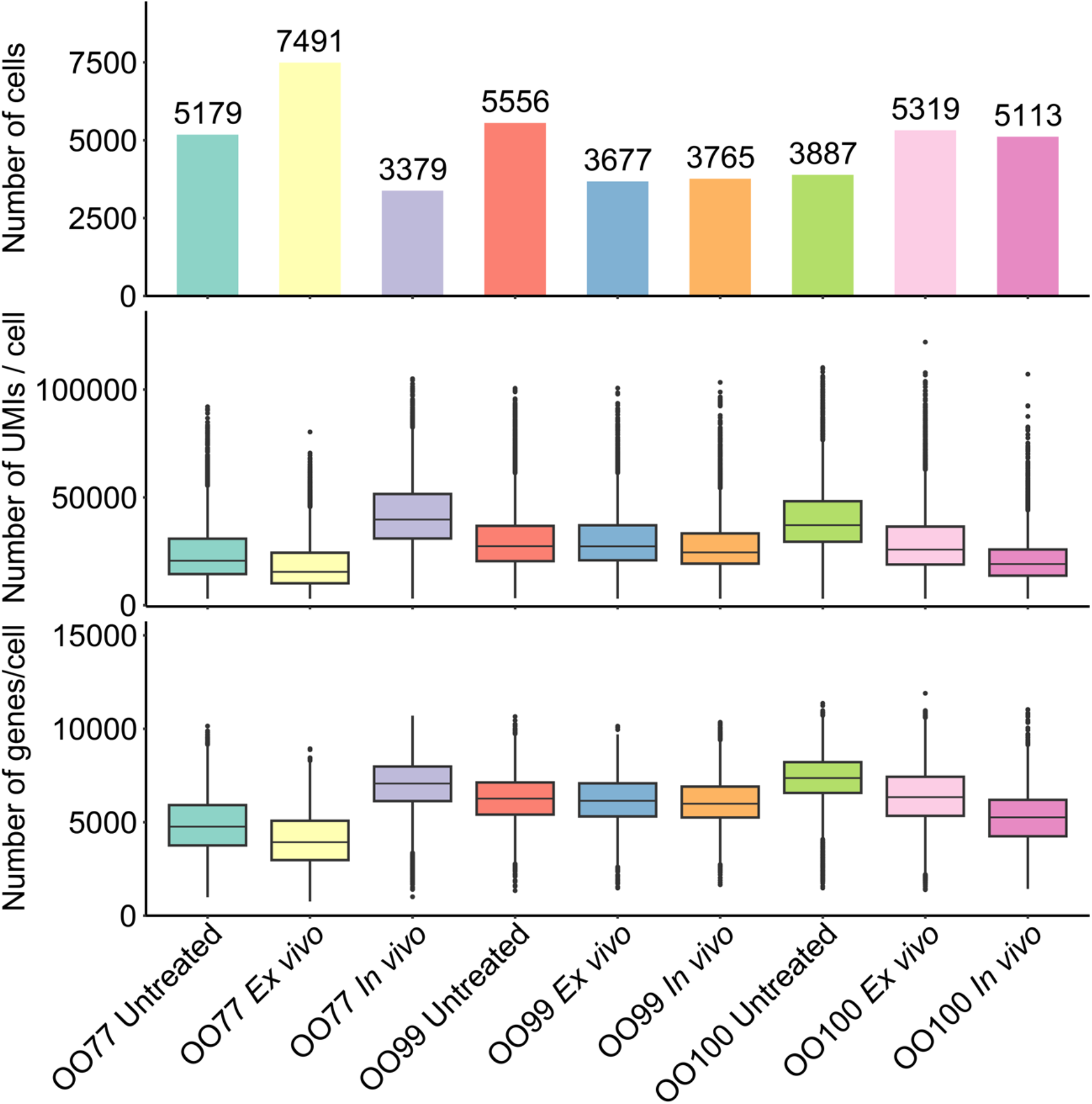
Single cell RNA sequencing of GEJ PDOs. Quality control of scRNA-seq data: Number of cells, number of UMIs per cell, and number of genes per cell in each sample.

**Supplementary Figure 2.**
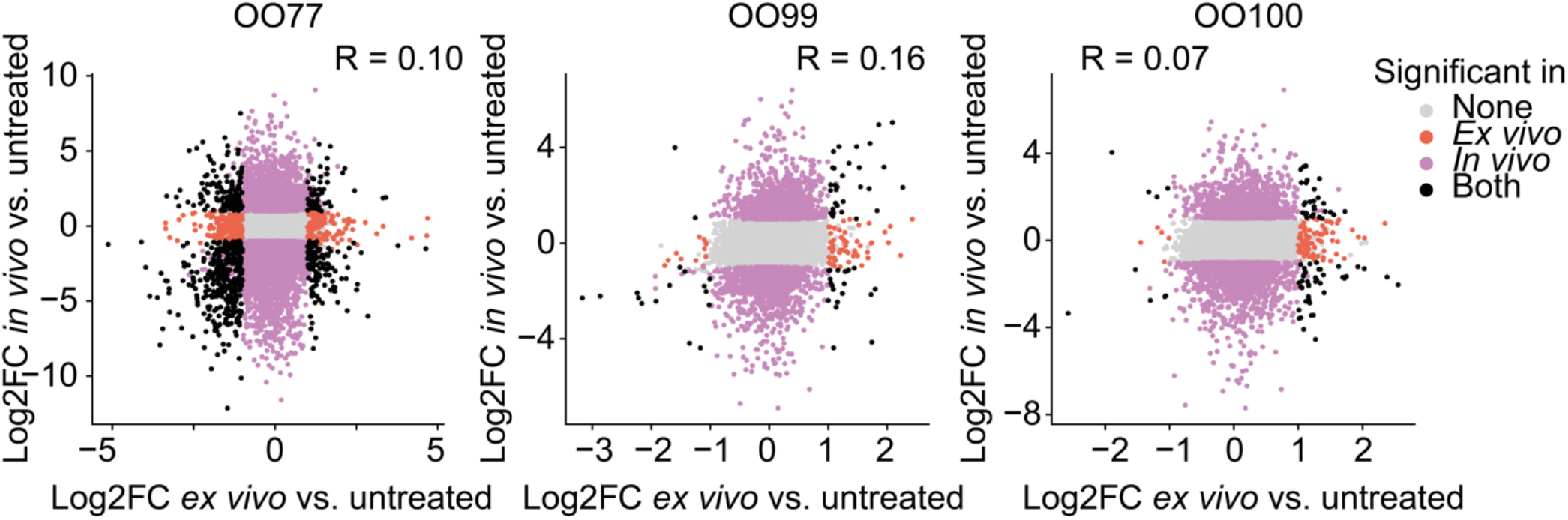
Pseudobulk differential analysis of post-treatment samples compared to the matched untreated samples. Scatter plot of the log2 fold change (log2fc) of gene expression. Each dot represents a gene, coloured by significance in the respective group with adjusted p-value < 0.01. Correlation statistics refer to Spearman’s R of the log2fc between both comparisons.

**Supplementary Figure 3.**
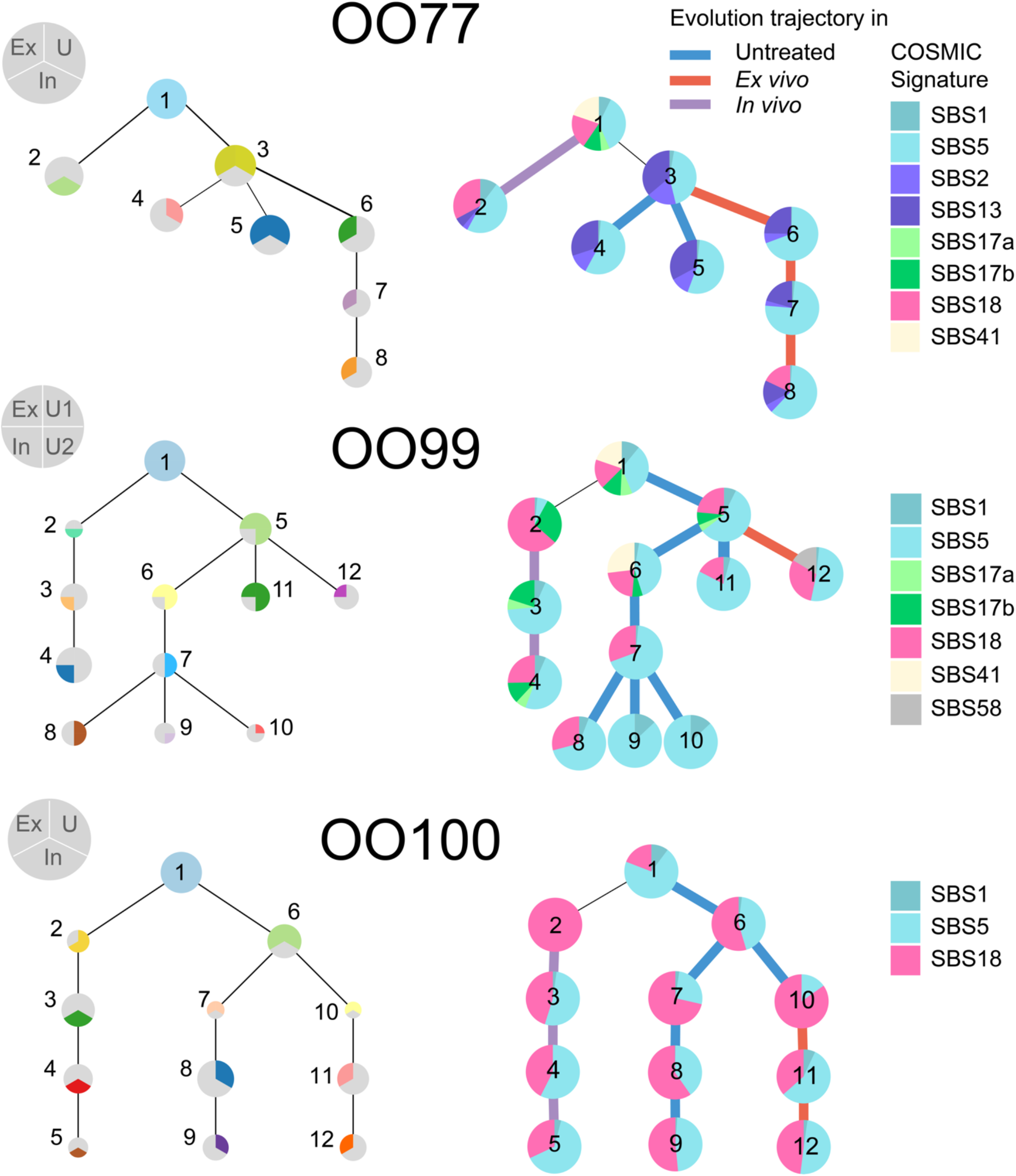
Bulk WGS-derived phylogenetic trees, tumor clone composition, and Single Base Substitution (SBS) mutational signatures. **Left:** Phylogenetic trees as output from Conipher. Each circle is a clone as a group of mutations which have the same cancer cell fraction. The fraction of the circle indicates in which samples the clone could be detected. Ex = *Ex vivo*, In = *In vivo*, U = Untreated. The area of the circle correlates with the number of mutations belonging to that clone. OO99 is reconstructed from two technical replicates. **Right**: SBS mutational signature composition of each mutation clone.

**Supplementary Figure 4.**
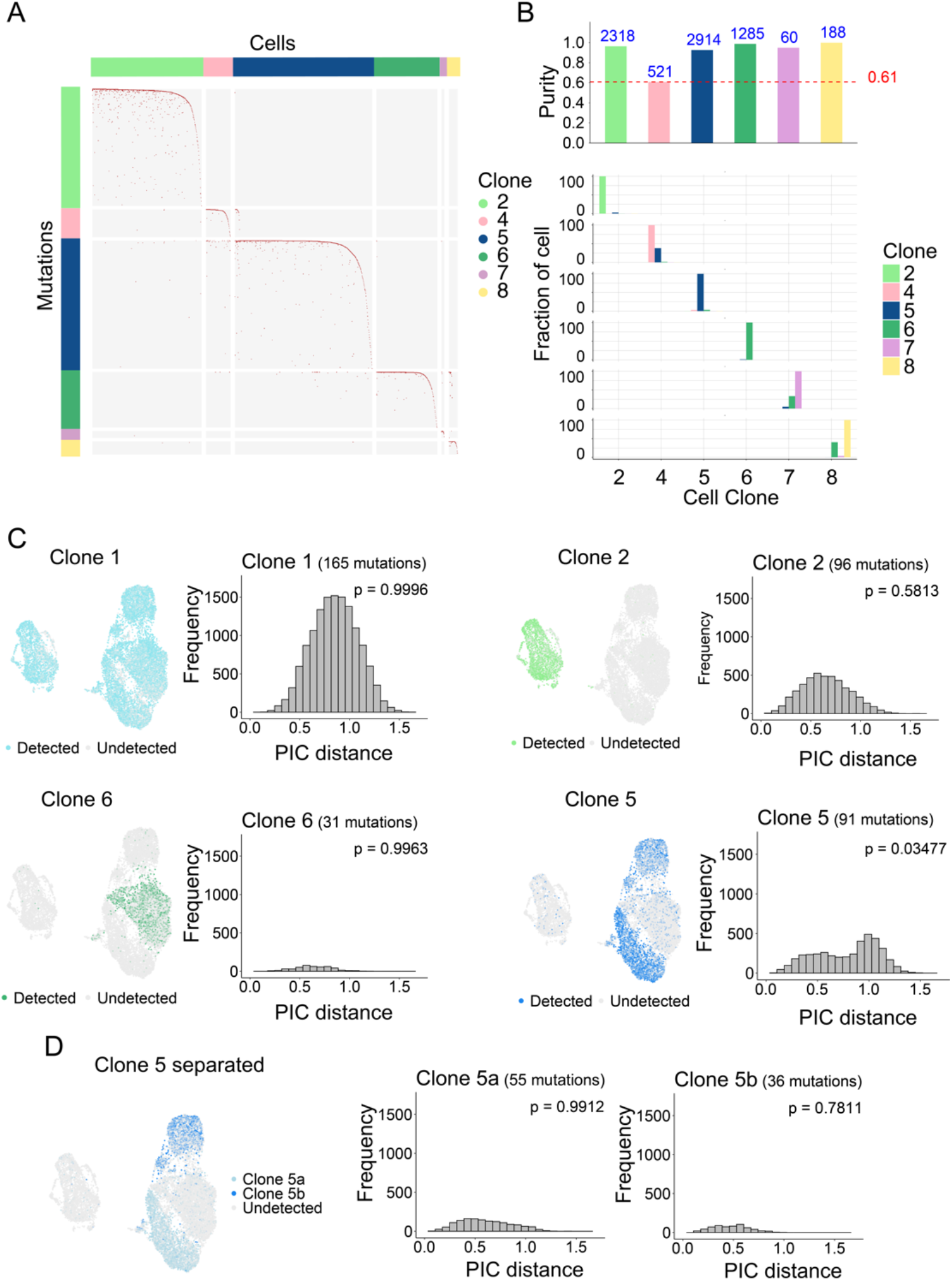
Reassessment of clonal reconstruction for OO77. (**A**) Mutation matrix with rows representing mutations from bulk WGS-derived phylogenetic trees, annotated according to their clone membership within the trees. Columns correspond to cells in scRNA-seq data. The heatmap was graphically enhanced using a gaussian smoothing strategy for better visibility. (**B**) **Top:** Purity score as percentage of cells in each cell clone that do not contain mutations from a separated branch in the bulk WGS-derived phylogenetic trees. The minimum purity of the respective reconstruction is marked in red. **Bottom:** Fraction of cells in each cell clone that contain mutations from each clone in the bulk WGS-derived phylogenetic trees. (**C**) Pairwise intra-clone distance for Clone 1, Clone 2, Clone 6, Clone 5. **Left:** UMAP representation of the cells’ single cell transcriptomes. Cells are coloured based on whether mutation of the specified bulk WGS tree clone is detected. **Right**: Histogram showing the pairwise intra-clone distances for the specified clones. P-values represent Hartigan’s Dip Test Statistics for Unimodality, testing the alternative hypothesis that the population exhibits more than one mode. (**D**) Result of the reconstruction separating Clone 5 into two subclones, 5a and 5b, as a consequence of the bimodality. **Left**: UMAP representation of the cells’ single cell transcriptomes. Cells are coloured based on whether mutations belonging to clone 5a or 5b are detected. **Right**: Histograms showing the pairwise intra-clone distances for clone 5a and 5b. P-values represent Hartigan’s Dip Test Statistics for Unimodality.

**Supplementary Figure 5.**
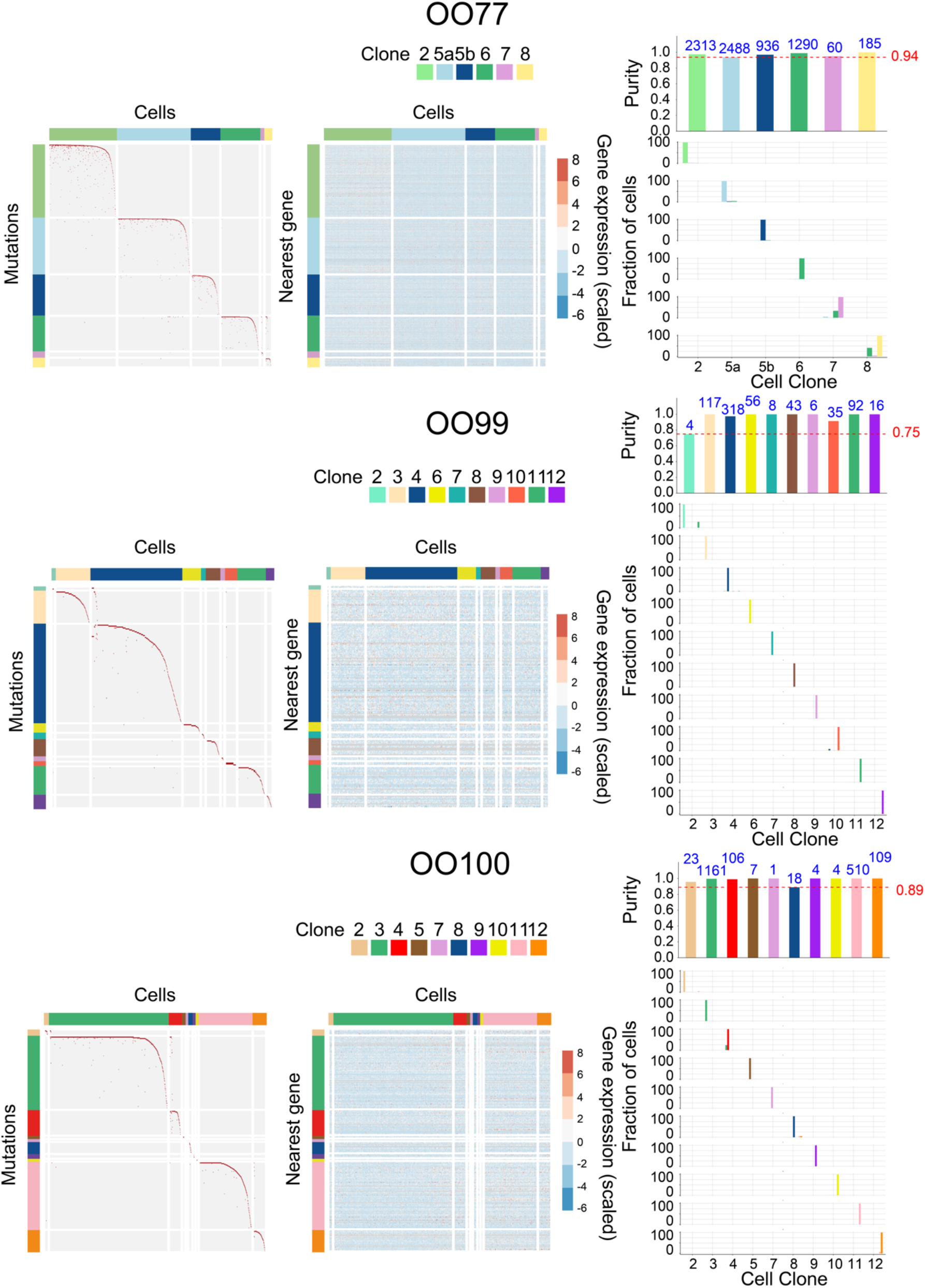
Co-occurance of mutations and cell clone purity in all three patients. **Left**: Mutation matrix with rows representing mutations from bulk WGS-derived phylogenetic trees, annotated according to their clone membership within the trees. Columns correspond to cells in scRNA-seq data and were annotated based on the final clone labelling. Heatmaps are graphically enhanced using a gaussian smoothing strategy for better visibility. **Middle**: Gene expression matrix. Rows are the nearest gene of the mutation in the Left panel and ordered to match the order of the corresponding mutation. Grey means that no gene could be assigned. **Right**: Purity score as percentage of cells in each cell clone that do not contain mutations from each clone in the bulk WGS-derived phylogenetic trees. The minimum purity of the respective reconstruction is marked in red.

**Supplementary Figure 6.**
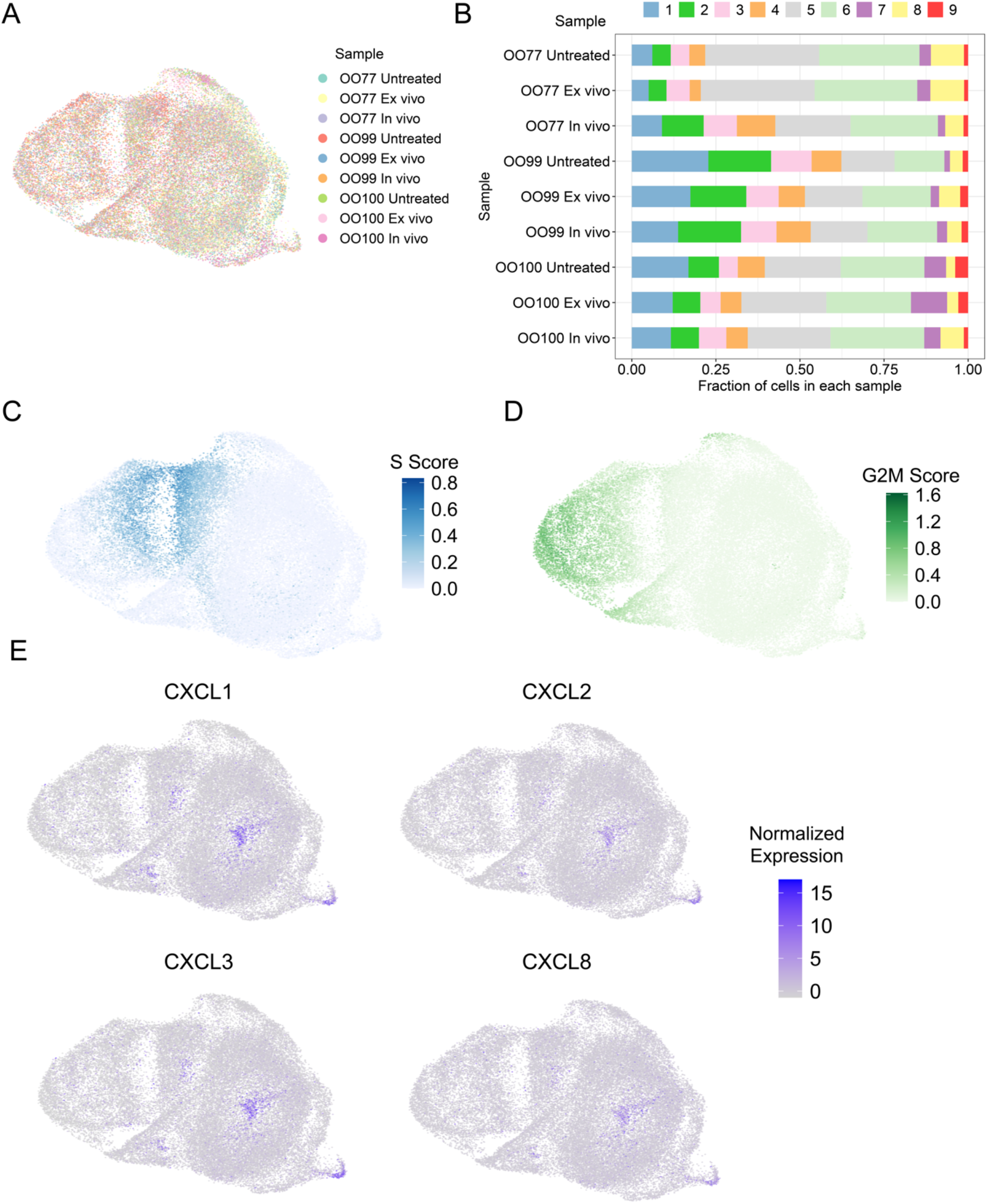
The samples share transcriptional cell states. (**A**) UMAP embedding of nine samples as in Figure 5B. Cells are colored based on sample of origin. (**B**) Cell distribution of the nine transcriptional clusters in each sample. (**C & D**) Cell cycle scores for each cell, computed based on the expression of S and G2/M phase genes, respectively, visualized relative to the UMAP of the single cell transcriptomes. (**E**) Expression of cytokines across all cells from organoid lines, visualized relative to the UMAP of the single cell transcriptomes.

## References

1. Park, J. Y. et al. Global lifetime estimates of expected and preventable gastric cancers across 185 countries. Nat Med (2025). https://pubmed.ncbi.nlm.nih.gov/40624406

2. Al-Batran, S. E. et al. Perioperative chemotherapy with fluorouracil plus leucovorin, oxaliplatin, and docetaxel versus fluorouracil or capecitabine plus cisplatin and epirubicin for locally advanced, resectable gastric or gastro-oesophageal junction adenocarcinoma (FLOT4): a randomised, phase 2/3 trial. Lancet 393, 1948–1957 (2019). https://pubmed.ncbi.nlm.nih.gov/30982686

3. Burrell, R. A., McGranahan, N., Bartek, J. & Swanton, C. The causes and consequences of genetic heterogeneity in cancer evolution. Nature 501, 338–345 (2013). https://pubmed.ncbi.nlm.nih.gov/24048066

4. Marusyk, A., Almendro, V. & Polyak, K. Intra-tumour heterogeneity: a looking glass for cancer. Nat Rev Cancer 12, 323–334 (2012). https://pubmed.ncbi.nlm.nih.gov/22513401

5. Schmäche, T. et al. Stratifying esophago-gastric cancer treatment using a patient-derived organoid-based threshold. Mol Cancer 23, 10 (2024). https://pubmed.ncbi.nlm.nih.gov/38200602

6. Krieger, T. G. et al. Single-cell analysis of patient-derived PDAC organoids reveals cell state heterogeneity and a conserved developmental hierarchy. Nat Commun 12, 5826 (2021). https://pubmed.ncbi.nlm.nih.gov/34611171

7. Seidlitz, T. et al. Human gastric cancer modelling using organoids. Gut 68, 207–217 (2019). https://pubmed.ncbi.nlm.nih.gov/29703791

8. Zhou, J. et al. Replication stress inducing ELF3 upregulation promotes BRCA1-deficient breast tumorigenesis in luminal progenitors. bioRxiv 2023.07.20.549893 (2023). https://www.biorxiv.org/content/10.1101/2023.07.20.549893.abstract

9. Zheng, H. et al. Targeted activation of ferroptosis in colorectal cancer via LGR4 targeting overcomes acquired drug resistance. Nat Cancer 5, 572–589 (2024). https://pubmed.ncbi.nlm.nih.gov/38291304

10. Kwon, O. H., Kim, J. H., Kim, S. Y. & Kim, Y. S. TWEAK/Fn14 signaling mediates gastric cancer cell resistance to 5-fluorouracil via NF-κB activation. Int J Oncol 44, 583–590 (2014). https://pubmed.ncbi.nlm.nih.gov/24337061

11. Hutti, J. E. et al. Oncogenic PI3K mutations lead to NF-κB-dependent cytokine expression following growth factor deprivation. Cancer Res 72, 3260–3269 (2012). https://pubmed.ncbi.nlm.nih.gov/22552288

12. Aibar, S. et al. SCENIC: single-cell regulatory network inference and clustering. Nat Methods 14, 1083–1086 (2017). https://pubmed.ncbi.nlm.nih.gov/28991892

13. Uetsuka, H. et al. Inhibition of inducible NF-κB activity reduces chemoresistance to 5-fluorouracil in human stomach cancer cell line. Experimental cell research 289, 27–35 (2003). https://www.sciencedirect.com/science/article/pii/S0014482703002234

14. Montagut, C. et al. Activation of nuclear factor-κ B is linked to resistance to neoadjuvant chemotherapy in breast cancer patients. Endocrine-related cancer 13, 607–616 (2006). https://erc.bioscientifica.com/view/journals/erc/13/2/0130607.xml

15. Wang, W., Cassidy, J., O’Brien, V., Ryan, K. M. & Collie-Duguid, E. Mechanistic and predictive profiling of 5-Fluorouracil resistance in human cancer cells. Cancer research 64, 8167–8176 (2004). https://aacrjournals.org/cancerres/article-abstract/64/22/8167/511974

16. Baldwin, A. S. Control of oncogenesis and cancer therapy resistance by the transcription factor NF-κB. The Journal of clinical investigation 107, 241–246 (2001). https://www.jci.org/articles/view/11991/files/pdf

17. Collins, J. L. & Kao, M.-S. The anticancer drug, cisplatin, increases the naturally occurring cell-mediated lysis of tumor cells. Cancer Immunology, Immunotherapy 29, 17–22 (1989). https://link.springer.com/article/10.1007/BF00199911

18. Stuart, T. et al. Comprehensive integration of single-cell data. cell 177, 1888–1902.e21 (2019). https://www.cell.com/cell/fulltext/S0092-8674(19)30559-8?dgcid=raven_jbs_etoc_email

19. McGinnis, C. S., Murrow, L. M. & Gartner, Z. J. DoubletFinder: doublet detection in single-cell RNA sequencing data using artificial nearest neighbors. Cell systems 8, 329–337.e4 (2019). https://www.cell.com/cell-systems/pdfExtended/S2405-4712(19)30073-0

20. Hafemeister, C. & Satija, R. Normalization and variance stabilization of single-cell RNA-seq data using regularized negative binomial regression. Genome biology 20, 296 (2019). https://link.springer.com/article/10.1186/s13059-019-1874-1

21. Robinson, M. D., McCarthy, D. J. & Smyth, G. K. edgeR: a Bioconductor package for differential expression analysis of digital gene expression data. Bioinformatics 26, 139–140 (2010). https://pubmed.ncbi.nlm.nih.gov/19910308

22. Ritchie, M. E. et al. limma powers differential expression analyses for RNA-sequencing and microarray studies. Nucleic Acids Res 43, e47 (2015). https://pubmed.ncbi.nlm.nih.gov/25605792

23. Hoffman, G. E. & Schadt, E. E. variancePartition: interpreting drivers of variation in complex gene expression studies. BMC Bioinformatics 17, 483 (2016). https://pubmed.ncbi.nlm.nih.gov/27884101

24. McLaren, W. et al. The Ensembl Variant Effect Predictor. Genome Biol 17, 122 (2016). https://pubmed.ncbi.nlm.nih.gov/27268795

25. Wang, K., Li, M. & Hakonarson, H. ANNOVAR: functional annotation of genetic variants from high-throughput sequencing data. Nucleic Acids Res 38, e164 (2010). https://pubmed.ncbi.nlm.nih.gov/20601685

26. Raine, K. M. et al. ascatNgs: Identifying Somatically Acquired Copy-Number Alterations from Whole-Genome Sequencing Data. Curr Protoc Bioinformatics 56, 15.9.1–15.9.17 (2016). https://pubmed.ncbi.nlm.nih.gov/27930809

27. Gurbich, T. A. & Ilinsky, V. V. ClassifyCNV: a tool for clinical annotation of copy-number variants. Sci Rep 10, 20375 (2020). https://pubmed.ncbi.nlm.nih.gov/33230148

28. Grigoriadis, K. et al. CONIPHER: a computational framework for scalable phylogenetic reconstruction with error correction. Nat Protoc 19, 159–183 (2024). https://pubmed.ncbi.nlm.nih.gov/38017136

29. Díaz-Gay, M. et al. Assigning mutational signatures to individual samples and individual somatic mutations with SigProfilerAssignment. Bioinformatics 39, btad756 (2023). https://pubmed.ncbi.nlm.nih.gov/38096571

30. Islam, S. M. A. et al. Uncovering novel mutational signatures by de novo extraction with SigProfilerExtractor. Cell Genom 2, None (2022). https://pubmed.ncbi.nlm.nih.gov/36388765

31. Frankell, A. M., Colliver, E., Mcgranahan, N. & Swanton, C. cloneMap: a R package to visualise clonal heterogeneity. bioRxiv 2022.07.26.501523 (2022). https://www.biorxiv.org/content/10.1101/2022.07.26.501523.abstract

32. Miller, C. A. et al. Visualizing tumor evolution with the fishplot package for R. BMC genomics 17, 880 (2016). https://link.springer.com/article/10.1186/s12864-016-3195-z

33. Goncearenco, A. et al. Exploring background mutational processes to decipher cancer genetic heterogeneity. Nucleic Acids Res 45, W514–W522 (2017). https://pubmed.ncbi.nlm.nih.gov/28472504

34. Tokheim, C. & Karchin, R. CHASMplus Reveals the Scope of Somatic Missense Mutations Driving Human Cancers. Cell Syst 9, 9–23.e8 (2019). https://pubmed.ncbi.nlm.nih.gov/31202631

35. Rogers, M. F., Shihab, H. A., Gaunt, T. R. & Campbell, C. CScape: a tool for predicting oncogenic single-point mutations in the cancer genome. Sci Rep 7, 11597 (2017). https://pubmed.ncbi.nlm.nih.gov/28912487

36. Mao, Y. et al. CanDrA: cancer-specific driver missense mutation annotation with optimized features. PLoS One 8, e77945 (2013). https://pubmed.ncbi.nlm.nih.gov/24205039

37. Tate, J. G. et al. COSMIC: the Catalogue Of Somatic Mutations In Cancer. Nucleic Acids Res 47, D941–D947 (2019). https://pubmed.ncbi.nlm.nih.gov/30371878

38. Gonzalez-Perez, A. et al. IntOGen-mutations identifies cancer drivers across tumor types. Nat Methods 10, 1081–1082 (2013). https://pubmed.ncbi.nlm.nih.gov/24037244

39. Chakravarty, D. et al. OncoKB: A Precision Oncology Knowledge Base. JCO Precis Oncol 2017, PO.17.00011 (2017). https://pubmed.ncbi.nlm.nih.gov/28890946

40. Lever, J., Zhao, E. Y., Grewal, J., Jones, M. R. & Jones, S. J. M. CancerMine: a literature-mined resource for drivers, oncogenes and tumor suppressors in cancer. Nat Methods 16, 505–507 (2019). https://pubmed.ncbi.nlm.nih.gov/31110280

41. Totoki, Y. et al. Multiancestry genomic and transcriptomic analysis of gastric cancer. Nat Genet 55, 581–594 (2023). https://pubmed.ncbi.nlm.nih.gov/36914835

42. Prashant, N. M. et al. SCReadCounts: estimation of cell-level SNVs expression from scRNA-seq data. BMC Genomics 22, 689 (2021). https://pubmed.ncbi.nlm.nih.gov/34551708

43. Muyas, F. et al. De novo detection of somatic mutations in high-throughput single-cell profiling data sets. Nat Biotechnol 42, 758–767 (2024). https://pubmed.ncbi.nlm.nih.gov/37414936

44. Love, M. I., Huber, W. & Anders, S. Moderated estimation of fold change and dispersion for RNA-seq data with DESeq2. Genome Biol 15, 550 (2014). https://pubmed.ncbi.nlm.nih.gov/25516281

45. Liberzon, A. et al. The Molecular Signatures Database (MSigDB) hallmark gene set collection. Cell Syst 1, 417–425 (2015). https://pubmed.ncbi.nlm.nih.gov/26771021

46. Kumar, N., Mishra, B., Athar, M. & Mukhtar, S. Inference of Gene Regulatory Network from Single-Cell Transcriptomic Data Using pySCENIC. Methods Mol Biol 2328, 171–182 (2021). https://pubmed.ncbi.nlm.nih.gov/34251625

47. Wingender, E., Dietze, P., Karas, H. & Knüppel, R. TRANSFAC: a database on transcription factors and their DNA binding sites. Nucleic Acids Res 24, 238–241 (1996). https://pubmed.ncbi.nlm.nih.gov/8594589

48. Hafemeister, C. & Satija, R. Normalization and variance stabilization of single-cell RNA-seq data using regularized negative binomial regression. Genome biology 20, 296 (2019). https://link.springer.com/article/10.1186/s13059-019-1874-1

49. The Cancer Genome Atlas consortium. Comprehensive molecular characterization of gastric adenocarcinoma. Nature 513, 202–209 (2014). https://pubmed.ncbi.nlm.nih.gov/25079317

